# Complex folding kinetics of multi-domain proteins disentangled with periodic forces by magnetic tweezers

**DOI:** 10.64898/2026.10.02.749805

**Authors:** César Augusto Quintana-Cataño, Andreas Hartmann, Michael Schlierf

## Abstract

Force spectroscopy enables direct investigation of the interplay between free energy and transition kinetics in proteins and nucleic acids. However, multi-domain proteins often display multiple intermediate structures with diverse stabilities and kinetics, making equilibrium conditions difficult to achieve and out-of-equilibrium experiments challenging to interpret. We introduce a single-molecule force spectroscopy assay using periodic forces and high-resolution magnetic tweezers to bridge different stabilities, selectively enhance intermediate populations, and rigorously analyze multi-domain protein folding kinetics at quasi-equilibrium. By applying well-defined periodic square force functions with tunable steady states, we first validate our framework on the two-state folder λ_6-85_, demonstrating that periodic force experiments can reproduce equilibrium kinetics. Extending this approach to the multi-domain protein mimic (Protein L)_8_, we show that tuning force, frequency, and amplitude enhances populations of distinct intermediate states. Finally, we dissect the entangled states of the hybrid two-domain protein λ-L, where a 100-fold difference in domain kinetics precludes equilibrium analysis, by expanding from a two-state to a three-state periodic force protocol to direct intermediate populations. Our assay and analysis framework open new possibilities for deciphering complex multi-domain folds and their multi-dimensional folding pathways.

## I. Introduction

Single-molecule force spectroscopy (SMFS) has become a widely used tool to assess protein and nucleic acid folding energy landscapes, quantify small molecule protein interactions and characterize the stabilities of molecular complexes [1–6]. Biomolecules are typically immobilized between a surface and a force probe, e.g. an AFM cantilever, an optical tweezers bead, a super-paramagnetic bead in magnetic tweezers, among many other possibilities. Upon exertion of force, structures and complexes are mechanically destabilized and at a characteristic force, change conformation, e.g. unfold in the case of protein folding. Direct access to kinetic rates of folding or unfolding reactions is provided via constant force experiments at equilibrium conditions, with advanced analysis models that connect the effects of force on transition rates and enable reconstruction of the underlying energy landscape. Current analysis models consider instrument properties, experimental design and elastic properties of studied molecules, which are often modeled as entropic springs, to extract transition barrier positions and heights [7–10]. If folding reactions are reversible and external conditions remain constant, measured work relates to free energy differences. Thus, comprehensive models on how force modulate kinetics can be developed even for complex processes of coupled folding and binding [11]. Unfortunately, constant force measurements often provide accessible reversible kinetics only in a very narrow range of forces due to an inherent relation between the force-dependent folding barrier and molecular size [12–14]. This limitation can render the determination of an unknown midpoint force (*F*_*m*_) challenging and laborious. Even more, multidomain proteins exhibit folding transitions at domain specific midpoint forces, potentially even with kinetics ranging from milliseconds to hours [15]. Hence, barely any force-spectroscopy studies of large soluble proteins at equilibrium are available.

Consequently, out-of-equilibrium SMFS, i.e. constant-velocity, force-ramp or force-jump experiments, has been the preferred method to study large complex molecules so far [16]. Noteworthy, in a typical force-ramp experiment with constant loading rate (*∂*_*t*_*F*), the unfolding force at which the contour length of a molecule suddenly changes is directly related to the mean unfolding rate ⟨*k*_f →u_⟩, without the need of reversible transitions [17,18]. Thus, out-of-equilibrium SMFS can resolve interconversion timescales that are inaccessible in equilibrium experiments, and, by extension, also obtain energy landscape parameters, which can be extracted for instance using fluctuation theorems [19–21]. However, the analysis and interpretation of out-of-equilibrium SMFS experiments requires extensive data acquisition, analysis time, and detailed physical modelling in order to gain accurate kinetic insights. For example, characterizing a single unfolding rate *k*_f →u_ requires hundreds of single-molecule transitions and for zero-force extrapolation even multiple experimental conditions covering a wide range of loading rates. This raises the question: Is it possible to combine the technical simplicity and high statistics of equilibrium data, with the out-of-equilibrium advantage to modulate and resolve kinetically decoupled transition rates?

In earlier studies, non-equilibrium steady states (NESS) were obtained using sinusoidal force functions [22–24], or zigzag force-ramps where suggested to investigate complex landscapes [25]. While such force spectroscopy protocols allowed access to short-lived states, a quantitative analysis of transition rates remained complex or impossible. In theory, NESS can be well described and complex kinetics modeled [26,27]. Here, we propose a combination of both approaches, equilibrium and non-equilibrium, in a quasi-equilibrium framework to extract kinetic information from NESS using periodic rectangular force functions. To this end, periodic forces between high and low values with defined dwell times are applied to a single or multidomain protein. A rectangular force spectroscopy protocol simplifies transition rate determination and provides flexibility to adjust force values and residence times, which opens the path to understand the intricate kinetics of complex molecules. Thereby, NESS are revealed, which were inaccessible before. We demonstrate our framework using a high resolution magnetic tweezers setup providing ultra-stable measurement conditions with nanometer-scale resolution [10,28,29]. Magnetic Tweezers enable stable force protocols due to their force-constraint geometry (i.e., negligible force drift from passive force-clamp conditions), thus, periodic forces for our quasi-equilibrium framework can be readily implemented. Starting with a model, we provide the theory for NESS analysis with an emphasis on intermediate-state enhancement by periodic forces. We demonstrate the validity of our approach by comparing measured equilibrium kinetics with NESS employing the fast-folding protein λ_6-85_. Subsequently, we study a multidomain protein consisting of the modular reference protein (Protein L)_8_ and illustrate the versatility of periodic forces and NESS with a Protein L - λ_6-85_ hybrid, where two proteins of similar size but different folding kinetics can be disentangled. Selective modulation of enhanced intermediate states is possible by careful considerations on the predicted kinetics. This framework will not only enable insights on how to investigate complex molecules under periodic forces, but also a clearer understanding of the behavior of multidomain proteins under periodic forces (or even cyclic) recurrent in biology.

## II. Modelling periodic forces in multidomain proteins

Periodic rectangular functions of force provide a simple, yet, comprehensive framework to investigate quasi-equilibrium states (Supplementary Note 1 for terminology). Periodic forces modulate a typical energy landscape of a multi-domain protein comprising in this picture a folded, an unfolded, and an intermediate state (Fig. 1). In the regimes of constant force, e.g., *F*(*t*) = *F*_low_ (Fig. 1, top), the equilibrium constant 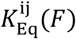 between state *i* and *j* is given by [1,30]:

**Figure 1.**
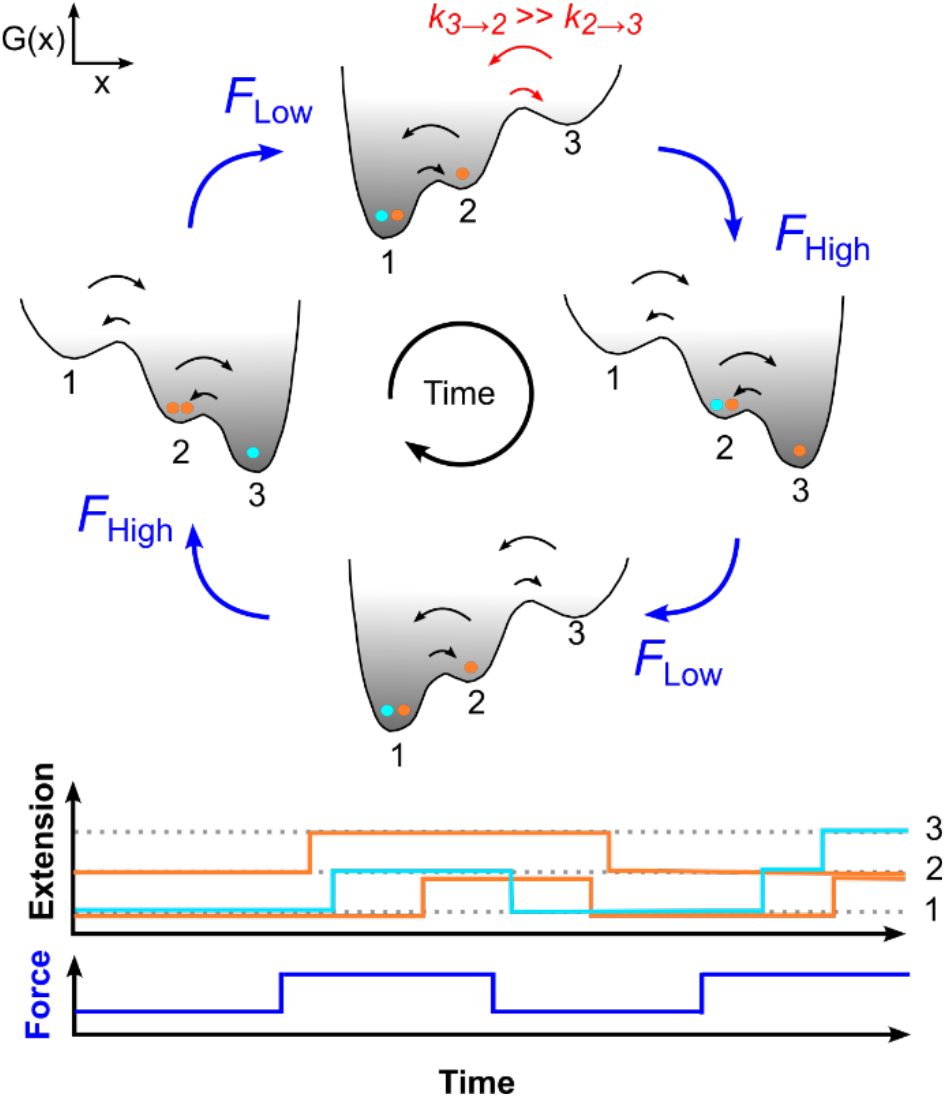
Effect of periodic forces on the one-dimensional folding landscape of a multi-domain protein. Schematics of a multi-domain protein comprising a folded state (1), an intermediate state (2), and an unfolded state (3). The energy landscape of the protein is tilted by forces *F*_high_ and *F*_low_, applied periodically. The shaded gray areas resemble equilibrium occupancies, with larger values (dark grey) at lower free energies. Under periodic forces the diffusion of individual proteins (cyan and orange dots) within the energy landscape is modulated leading to an enhanced intermediate state occupancy. This force dependent modulation of the kinetic rates can be modeled, for instance with *Monte Carlo* simulations (exemplary schematics down with each state having a different extension), with kinetics extracted from force spectroscopy models.

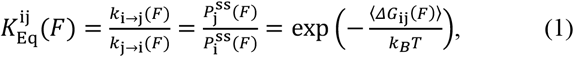

where the steady state occupancies of the folded state 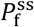 would have the highest probability at low force. Here, *k*_i→j_ denotes the transition rate from state i to state 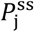 the steady state probability of finding the molecule at state j, and Δ*G*_ij_(*F*) ≡ *G*_i_ (*F*) − *G*_j_(*F*), the force-modulated free energy difference between state i and j. In this formalism, the effects of force on kinetics was derived following Kramers’ rate theory [18,31,32],

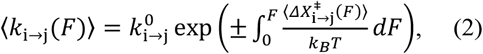

with 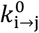 and 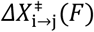 being the transition rate at zero force and the force dependent distance to the barrier associated to the transition between state *i* and *j*, respectively. When translating Kramers’ barriers into detailed balance (see Supplementary Note 3), it is important to distinguish between minimum and averaged free energies. To account for periodic forces, and thus out-of-equilibrium conditions, eq. 2 has to be implemented such that *k*_i →j_ (*F*) changes over time with *F*(*t*). We imagine a periodic force experiment (Fig. 1), where each of the transition rates oscillates between two states, i.e., *k*_i →j_ (*F*_low_) and *k*_i →j_(*F*_high_). To show the difference in occupancy resulting from out-of-equilibrium kinetics, the conformational diffusion of three individual proteins (Fig. 1, cyan and orange dots) is illustrated within the energy landscape showing, e.g., two folded and one intermediate state molecule (Fig. 1, top). Once the force shifts from *F*_low_ to *F*_high_, most of the proteins would equilibrate in an unfolded state. However, if the cycle shifts the force back to *F*_low_ before the system equilibrates, the transition to the unfolded state is quenched mid-way leading to a higher occupancy of molecules in the intermediate state (steady-state occupancies). In contrast to a fully equilibrated system, where either the unfolded or the folded states would dominate (equilibrium occupancies).

Therefore, the effect of force on stability and kinetics can be described by considering Bell-Evans’ force dependent barriers, Kramers’ conformational diffusion and physical models including polymer spacers and instrumentation (Supplementary Note 2) [6,11,32–34]. The resulting force dependent kinetic rates *k*_i →j_ (*t*) allow the prediction of the time dependent state occupancies *P*_i_(*t*) under the effect of an external periodic force *F*(*t*) using the following master equation [35],

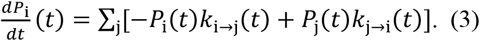

For a periodic force alternating between two distinct forces, *F*_low_ and *F*_high_ (Fig. 1), the transition rates oscillate between two discrete set of values, 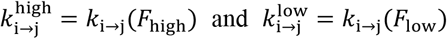. At equilibrium, deducing steady states is straightforward by means of eigenvectors of the transition probability matrix that describes the evolution of the system (Supplementary Note 4). Out of equilibrium, we start from the initial values *P*_i_ (*t* = o), e.g. at *F*_low_, and maintain border conditions between force transitions to ensure continuity of *P*_i_ (*t*). The time dependent state occupancies *P*_i_ (*t*) of complex Markov models are reproduced by *Monte Carlo* simulations and numerical solutions of the master equation allowing to deduce steady states and residence time distributions. Thus, in the following paragraphs we use periodic forces to study a simple two-state folder, a multi-domain protein and dissect kinetically entangled two-domain proteins.

## III. A two-state folder under periodic forces

In order to demonstrate the quasi-equilibrium framework for a small globular protein, we applied periodic forces to λ_6-85_, a five helix bundle fragment of the Lambda repressor, a protein folding model with microseconds-scale downhill folding [36]. Previously, we showed the effects of constant forces on the kinetics of λ_6-85_ using a custom-built high resolution magnetic tweezers setup [10]. Here, we study the mechanics of λ_YA_, a variant of λ_6-85,_ with a folding time of ~20 µs at its thermal denaturation midpoint [37]. To investigate λ_YA_ in magnetic tweezers, we embedded the protein between an N-terminal HaloTag for covalent surface immobilization and two C-terminal Spy0128 domains as a stiff spacer followed by an AviTag for *in vitro* biotinylation and subsequent binding to a magnetic bead [13,38] (Fig. 2a). λ_YA_ showed two-state folding in the magnetic tweezers and we studied in a first set of experiments unfolding and refolding kinetics at constant forces to obtain an equilibrium folding kinetics reference data set before switching to periodic forces. We measured hundreds of unfolding and refolding transitions of nine different λ_YA_ proteins at six different constant forces in the range ~7-10 pN (Fig. 2b). The observed unfolding-refolding transitions showed the expected λ_YA_ extension and contour length change allowing us to identify a fingerprint for the two-state transition. We extracted from the data set state-occupancy histograms at each force 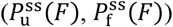, and using a thresholding algorithm, residence times in the unfolded or folded state and, by extension, the respective folding *k*_u →f_(*F*) or unfolding *k*_f →u_(*F*) rates (Fig. 2b and c). From the average ⟨*k*_f →u_(*F*)⟩ and ⟨*k*_u →f_(*F*)⟩ we estimated zero-force transition rates ⟨*k*_i →j_ (*F* = 0)⟩ and the distance of each state to the transition state using different extrapolation models (Supplementary Note 5) [10]. We found good agreement with earlier optical tweezers measurements on λ_YA_ [39]. Subsequently, we performed periodic force experiments on the same, well-characterized construct. With periodic forces, we varied four parameters: the high force *F*_high_ and corresponding time *T*_high_; and the low force *F*_low_ and corresponding time *T*_low_ (Fig. 2d), thus we could push λ_YA_ into different conformational NESS. In particular, we tested the effects on λ_YA_ (un)folding for different variations of the periodic force-protocols, with varying period (*T*_high_ + *T*_low_), amplitude (*F*_high_ − *F*_low_), or a single parameter at a time. Using the models extracted from equilibrium data (Supplementary Note 4), we could predict the steady state stabilities of λ_YA_ (characterized by 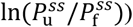 under periodic forces. For this we plot 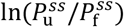 as quantified from the histograms of extensions, with respect to expected values from *Monte Carlo* simulations (Fig. 2e and Supplementary Note 6) (or alternatively using the master equation solution). We found excellent agreement with deviations <*k*_*B*_*T*/2 (Fig. 2f). We further validated this approach using *in silico* Langevin dynamics simulations mimicking both conditions, equilibrium and NESS (Fig. 2f). Variations in the parameters of a given periodic force protocol such as amplitude or frequency have effects on steady state occupancies, sensitive to kinetics and sometimes counterintuitive. For instance, due to the different periodic forces, lifetime distributions of the same state become multi-exponential rather than single-exponential in a equilibrium experiment. This can be taken into account during analysis. Interestingly at distinct periodicities and forces, we could drive λ_YA_ also into stochastic resonances as described earlier for nucleic acids or Talin (Supplementary Note 7) [24,40].

**Figure 2.**
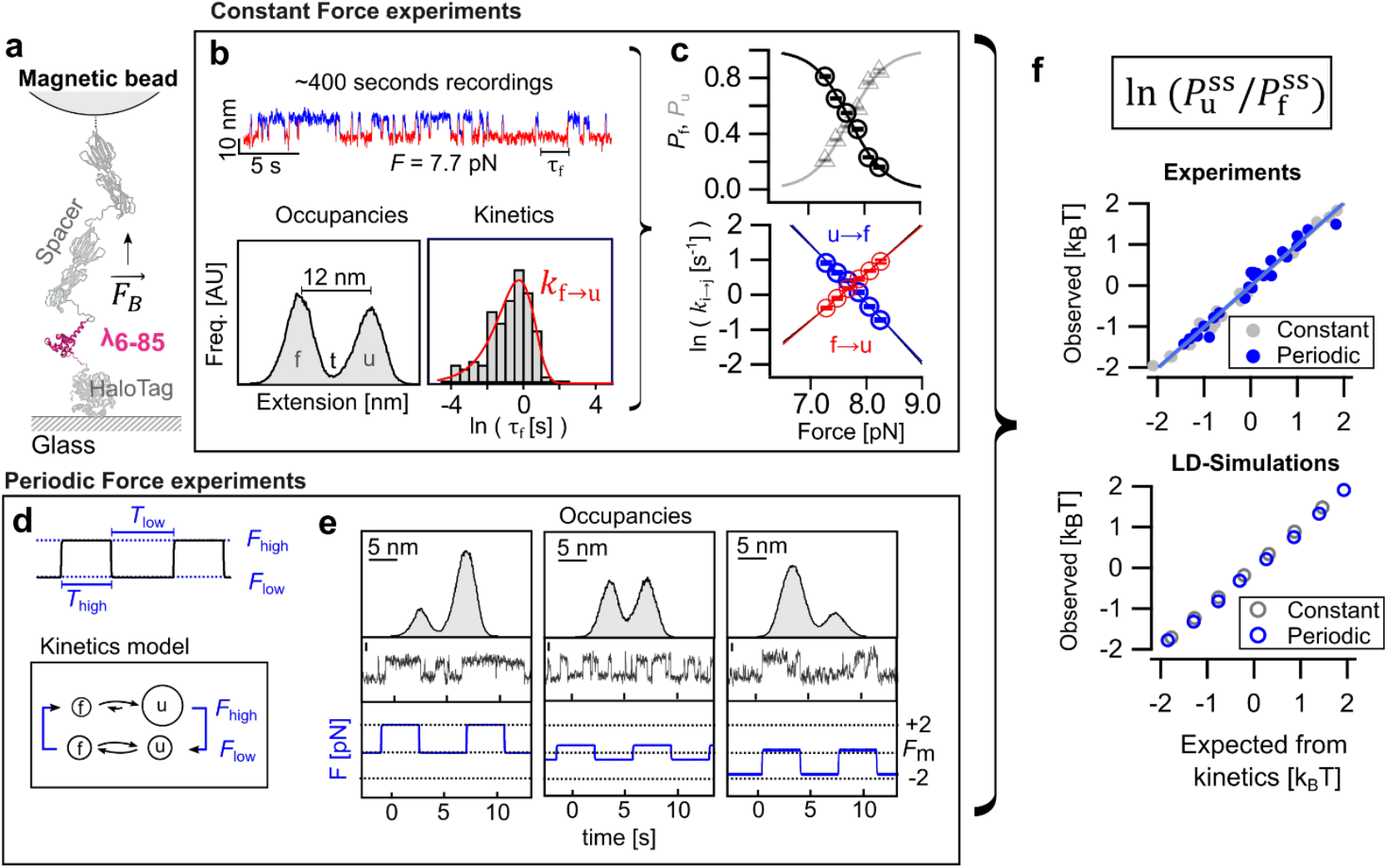
λ_YA_ stability under periodic forces agrees with predictions from two-state kinetic models. (a) Experimental design overview. **Constant Force Panel** (b) λ_YA_ (un)folding kinetics measured by magnetic tweezers [10]. Two-state behavior (folded f, unfolded u) characterized at multiple constant forces and state occupancies extracted by double-Gaussian fitting of extension histograms assuming P_f_ = A_f_ / (A_u_ + A_f_). Transition rates were determined using single-exponential fits to dwell-time distributions (red: folded state; blue: unfolded state). (c) P_f_, P_u_, *k*_f→u_, and *k*_u→f_ as a function of force (solid lines; see Supplementary Note 2). P_f_ and P_u_ fitted with a sigmoidal phenomenological curve. *k*_*i* →*j*_ described by kinetic extrapolation models [10]. **Periodic Force Panel** (d) Square-wave force protocols applied following constant-force characterization. Kinetic model uses two force-dependent sets of transition rates: *k*_i →j_ (*F*_high_) and *k*_i →j_ (*F*_low_). (e) Three representative square-wave protocols with corresponding extension vs. time traces. Scale bars: 5 nm; traces shown as zoomed excerpts from full hundreds of seconds recordings. (f) Methods validation via *Monte Carlo* simulations predicting non-equilibrium steady-state stability 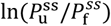. Input rates drawn from: (i) experimental constant-force data, or (ii) constant-force Langevin dynamics simulations. Linear regressions show excellent agreement between simulated and observed stabilities.

## IV. Multi-domain modular proteins under periodic forces

After confirming the analysis and kinetic models on a simple two-state folder protein, we expanded the experiments to a multi-domain protein construct with independent domains, the modular recombinant protein (Protein L)_x8,_ hereafter L_x8_ for simplicity. L_x8_ is composed of 8 identical repeats of the widely studied B1 sub-domain of Protein L [28,41] flanked by an N-terminal HaloTag and a C-terminal biotin (Fig. 3a). Earlier studies have reported a midpoint force of *F*_*m*_~8 pN and monomer (un)folding rates of *k*_u →f_(*F*_*m*_)~o.oo4 s^−1^ [42]; at forces above ~10 pN, only unfolding is measurable as the folding rate is one order of magnitude lower than the unfolding rate. At low forces below ~4 pN refolding dominates over unfolding, now with folding rates larger by roughly two orders of magnitude [42]. Having in mind that at low and high forces we can neglect contributions from either unfolding or folding (respectively), we decided to perform the periodic forces between 1 pN and 40 pN, for the low and high force, subjecting L_x8_ to at least 150 cycles with different periods (Fig. 3b). L_x8_ unfolded in several seconds at 40 pN (Fig. 3b inset i) and each of the eight domains gave rise in stepwise extensions with a characteristic difference in contour length of Δ*L*_fu_ *≈* 18.6 *nm* [41]. We assigned states by the number of unfolded domains - *N*_u_ from 0 to 8 -, each state with a characteristic extension at a given force (Fig. 3b). Polyproteins cover a wide range of timescales due to state degeneracy: at 40 pN, the first unfolding step of L_x8_ happened from the unfolding of any of the eight folded repeats (degenerate transition) with an average rate *k*_0→1_, while the last unfolding step (*k*_7→8_) resembled monomer unfolding kinetics, hence *k*_0→1_ > *k*_1→2_ > (…) > *k*_7→8_ [43]. Within these estimations we expect unfolding rates on the order of *k*_7→8_(4o pN)~o.3 s^−1^ [41] and *k*_0→1_(4o pN)~(*N*_u_ = 8) · o.3 s^−1^ = 2.4 s^−1^, thus spanning approximately an order of magnitude. In consequence, depending on the residence time at the high force, we can likely unfold also only a part of the multi-domain protein and thus enhance the occupancy of intermediate states. At 1 pN, we cannot directly observe protein folding due to the limited signal-to-noise, though through switching to the high force afterwards, allows for an indirect readout of the folding state of the multi-domain protein. Similarly, at shorter residence time in the low force regime, only a fraction of the domains will be able to refold.

**Figure 3.**
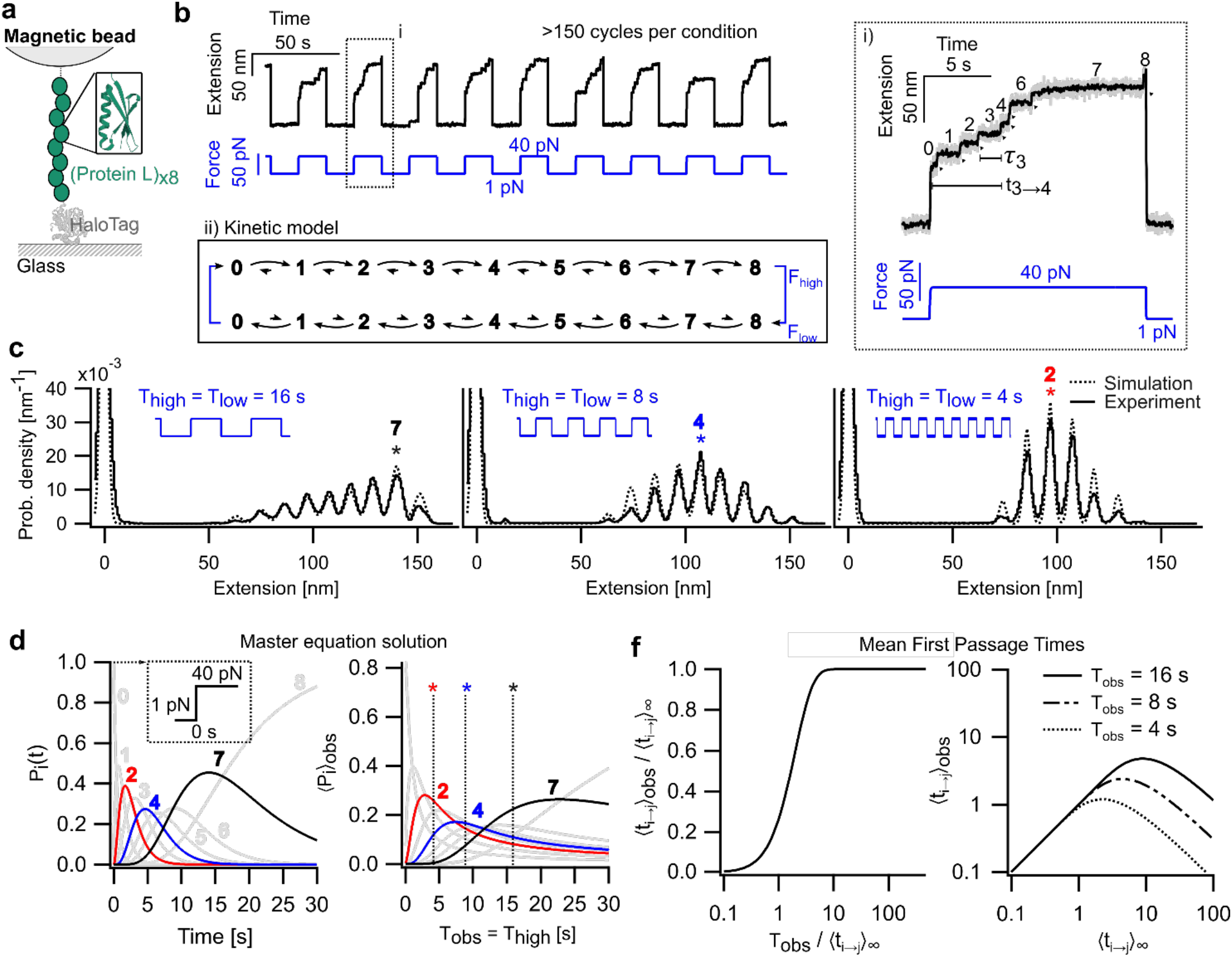
Periodic forces enhance the occupancy of intermediate states during unfolding of a modular reference protein. (a) Experimental design schematic. (b) Representative periodic force protocol: *F*_high_ = 40 pN, *F*_low_= 1 pN, *T*_high_ = 16 s, *T*_low_ = 16 s. More than 150 cycles collected per experimental condition. Inset (i): Up to 8 unfolding steps observed for Lx8, reflecting multiple degenerate states; dwell times (e.g., state 3) and first passage times (e.g., *t*_3→4_) indicated. Inset (ii): Proposed kinetic model derived from dwell times and first passage times. (c) Extension histograms at three different periods (*T*_high_ values). Different unfolding states of Lx8 are enhanced at each period; the state with highest occupancy is marked with an asterisk (*). Overlaid *Monte Carlo* simulations (rate values from *T*_high_ = *T*_low_ = 16 s; Supplementary Note 8) agree with experimental outcomes. (d) Numerical solution of the master equation. State occupancy *P*_i_(*t*) as a function of time; average occupancy up to observation time *T*_obs_, denoted ⟨*P*_i_⟩_obs_, is shown for each state. Experimental *T*_high_ values (corresponding to panels in c) are indicated. Evolution of enhanced states 2, 4, and 7 is highlighted; the highest ⟨*P*_i_⟩_obs_ at a given *T*_high_ identifies the most populated state. (f) Analytical equations for the effect of finite observation time on first passage times under periodic forces, providing a systematic framework for interpreting the data.

While changing periods, different intermediate states where enhanced, which is readily visible on the extension histograms extracted from ~ 150 cycles each (Fig. 3c). With long periods of 32 seconds, every folding state in the unfolding landscape of L_x8_ was sampled with *N*_u_ = 7 being the most populated state. While at short periods of 8 seconds, we observed up to state *N*_u_ = 5, with *N*_u_ = 2 the most populated (Fig. 3c). Thus, we wondered if we could rationalize which intermediate state is enhanced for a given periodic force protocol? As for λ_YA_ we computed the expected evolution of states with both the master equation and *Monte Carlo* simulations. Yet, we required individual folding and unfolding kinetics of single protein L domains, and thus, using the measurement with a period of 32 seconds, we determined the dwell time of each state (e.g. *τ*_3_ for state 3 in Figure 3 b); and the cumulative time from the moment we apply a high force (e.g. *τ*_3→4_ in Figure 3 b), or in other words the first passage time from state 3 to state 4. In agreement to theory, rates scale with degeneracy plus so-called linker effects due to conformational restrictions the more domains are already unfolded. Taking into account these transition rates, we readily reproduced the experimental occupancies with *Monte Carlo* simulations also as a function of the period of applied forces (Fig. 3c). The occupancy of each state was also tracked with a master equation, comparing the observed values with time averaged occupancies. The probability evolution of each unfolded state after applying F = 40 pN show that at observation times *T*_high_ = 16 *s*, 8 *s* and 4 *s*, the enhanced states are states *N*_u_ = 7, 4 and 2 respectively (Fig. 3d). Notably, the rectangular force modulation allows, in contrast to continious force modulations, for a simple analytical derivation of apparent first passage times (*t*_i →j_ *cf*. Fig. 3b), and their corresponding time-averaged values accounting for a finite observation time due to a force-quench at a time *T*_obs_ (see Supplementary Note 9):

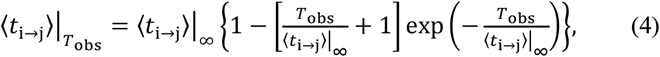

where the observation time *T*_obs_ corresponded to *T*_high_ in the used periodic-force protocols, ⟨*t*_i →j_⟩_∞_ the mean first passage time when the observation time is infinite, and 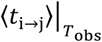 when its finite. If the observation time *T*_obs_ is sufficiently long 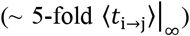, then 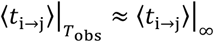. If the observation time *T*_obs_ ~ ⟨*t*_i →j_⟩_∞_, the slowest transitions will be truncated. We note that state occupancies are hard to describe analytically [43], yet, 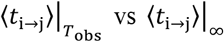 already resembles the shape of the measured extensions histograms. To transfer the mean passage times (eq. 4) from the context of periodic forces to cyclic forces, which are relevant for mechano-chemistry [44–46], *T*_obs_ must be drawn from a probability distribution instead of a constant value, for instance a Poisson process that simulates the cyclic action of a molecular motor.

Taken together, using a rigorous analysis of periodic force experiments with *Monte Carlo* simulations and analytical mean passage time descriptions, kinetics of modular proteins with independent, non-interacting domains where force is equally applied to each domain separately can be well described. Thus, we provide means to estimate for a given periodic force protocol, which intermediate states will be enhanced and a analytical solution for the effects of periodic forces in first-passage times. Notably, if multi-domain proteins do not follow such a description, an interaction between the domains is likely. Now with these tools in hand, we showed ways to disentangle also complex multi-domain proteins that would otherwise be hard to investigate.

## V. Dissection of entangled states with periodic forces

Molecules with multiple domains often present entangled states, e.g. either due to very similar equilibrium end-to-end distances or due to large discrepancies in kinetics, which are hard to resolve. We considered in this case a model protein consisting of a λ_YA_ domain and a Protein L domain in series (hereafter λ_YA_-L, Fig. 4a). Measured separately, the folding rates of both domains differ by two orders of magnitude at their respective midpoint force, 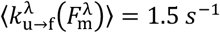 and

**Figure 4.**
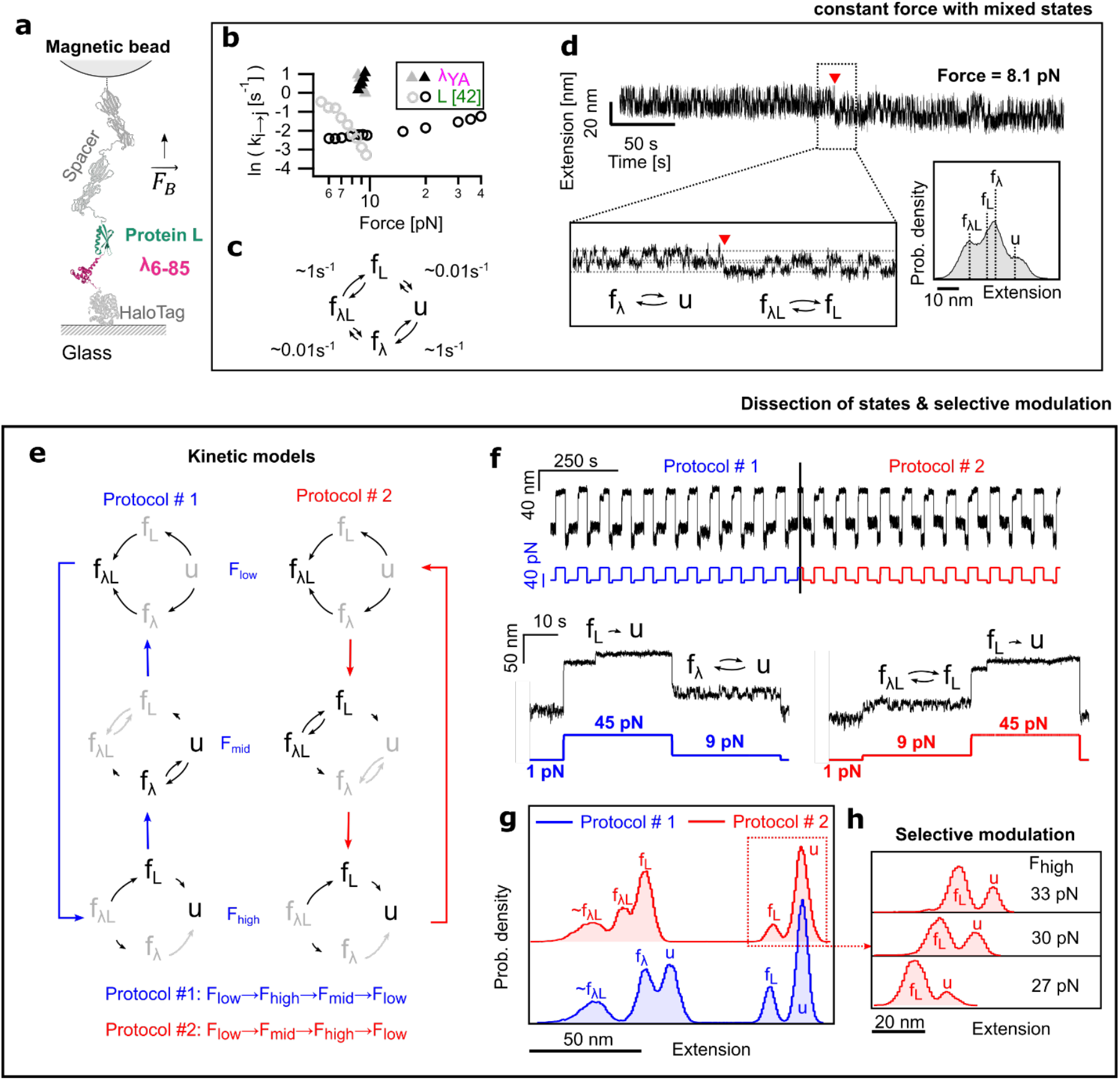
Kinetic dissection and selective modulation of entangled states with periodic forces. (a) Schematics of the hybrid λ_YA_-L construct, comprising two domains of similar size with distinct folding/unfolding kinetics. (b) Measured folding and unfolding rates for λ_YA_ and monomeric L tethers as a function of force (monomeric L kinetics adapted from [42]), illustrating differences in timescale and force dependence. (c) Kinetics of λ_YA_-L near the midpoint force (*F*_m_), revealing four distinct states; end-to-end extensions are entangled owing to comparable subdomain sizes. (d) Representative single-molecule trace exhibiting distinct kinetics for λ_YA_ and L domains but overlapping extensions, the red arrow points at L folding; inset histogram shows three observed states rather than the expected four. (e) Kinetic models used to dissect states with two different periodic force protocols; applying low (1 pN), middle (near λ_YA_ midpoint), and high (45 pN) forces. Two force-cycle protocols are shown, differing in the sequence of applied forces. (f) Periodic force measurements for both protocols, with a zoom-in to an individual cycle. (g) Extension histograms for both cycles, resolving five states. (h) Selective modulation of the two largest extension states as by means of a changing *F*_high_ while keeping all other parameters constant.

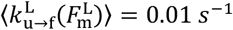 (Fig. 4b). Due to different *F*_*m*_s and stark differences in kinetics, reversible transitions for both domains are hard to observe at a single constant force (Fig. 4c and d), a scenario, which is likely for many large multidomain proteins. Besides transition rates, many protein domains are often on a similar size, offering the additional challenge that similar extension changes are to be observed for both domains at equilibrium. Ideally for λ_YA_-L, at constant force we expect four states: both domains folded (*F*_*λ*L_), only λ_YA_ folded (*F*_*λ*_), only L folded (*F*_L_), and both domains unfolded (u) (Fig. 4c and d). Using the end-to-end extension of state f_*λ*L_ as a zero-reference, for each state we expect in principle the following extensions: *X*_MLN_(*F*_*λ*L_, 8*pN*) = o, *X*_MLN_(*F*_*λ*_, 8*pN*) = 8.1 *nm, X*_MLN_(*F*_L_, 8*pN*) = 11.6 *nm* and *X*_MLN_(*F*_*λ*L_, 8*pN*) = 19.7 *nm*, with only a 3 nm difference between states f_L_ and f_*λ*_. Measured extensions from λ_YA_-L show individual populations with FWHM of ~5 − 10 *nm*, and hence states f_L_ and f_*λ*_ merge, such that an extension histogram shows only three apparent peaks (Fig. 4d, inset). Out of equilibrium conditions are often used to solve these challenges driving molecules to conditions at which states can be separated [22], yet, due to non-linearities, these are hard, if not impossible, to analyze.

Periodic forces as introduced above are NESS experiments and thus can be used to dissect states that are otherwise hard to differentiate, and even selectively modulate their occupancies. Noteworthy, λ_YA_ and L have very different mechanical properties, meaning they react to external force differently. For example, the distance from the folded state to the transition state differs nearly by an order of magnitude with 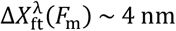 for λ_YA_ and 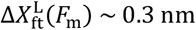 for protein L [13]. Since this distance can be seen as a leverarm for the force transmission, the force-dependent increase in the unfolding rate for protein L is much less than for λ_YA_; a property we can exploit to dissect both domains with periodic forces. To this end we used protocols oscillating among three different forces *F*_low_, *F*_mid_ and *F*_high_ (Fig. 4e and f), each with a characteristic observation time of *T*_low_, *T*_mid_, and *T*_high_. Consider a protocol of the form *F*_low_ → *F*_high_ → *F*_mid_ → *F*_low_, with a proposed kinetic model as schematized in Figure 4e. *F*_low_, *F*_mid_ and *F*_high_ and their corresponding observation times *T*_low_, *T*_mid_, and *T*_high_ were selected such that transition rates of interest are measurable within their observation time windows, while other rates become negligible. To dissect states of λ_YA_-L we proceeded then as follows: We start from *F*_low_ = 1 *pN* (*T*_low_ = 1o *s*), with a likely fully refolded protein in the state *F*_*λ*L_. Then jump to *F*_high_ = 45 pN (*T*_high_ = 3o *s*), where we expect both domains to unfold, though λ_YA_ unfolds very fast, hard or impossible to identify, thus we predominantly observed *F*_L_ → *U* transitions. Subsequently, the force is reduced to 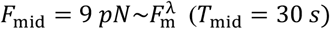, showing reversible transitions corresponding here to λ_YA_ (*F*_*λ*_ ⇆ u). In principle L could refold as well, yet due to the separation of timescales by more than two orders of magnitude 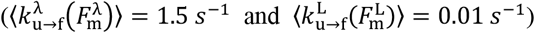, we predominantly will observed Λ_ya_ reversible (un)folding transitions. We note, that the distinct states were separated not only by kinetics, but also by means of their corresponding force-extension response, a distinct, characteristic of each protein domain. Using such a periodic force protocol, the resulting extension histogram shows five distinct peaks (Fig. 4g, protocol #1), which can be assigned as follows: (i) one for entangled states at *F*_low_ (which is dominated by f_*λ*L_), and (ii) four that can be traced back to unique states at *F*_mid_ and *F*_high_. Interestingly, the order of the periodic force cycle remains crucial. Note the occupancy differences between the extension histograms for the protocol #1 (order *F*_low_ → *F*_high_ → *F*_mid_ → *F*_low_) vs for the protocol #2 (order *F*_low_ → *F*_mid_ → *F*_high_ → *F*_low_), enhancing different transitions and emphasizing that periodic forces are NESS experiments (Fig. 4g). These changes in occupancies are also affected by linker effects on folding rates [10]. For both cycle types, the lifetimes of each individual state can be readily extracted using automatic thresholding and due to the discrete forces, the lifetimes can be readily converted to transition kinetics (Supplementary Note 10). In a sinusoidal or other force pattern, such lifetimes would not be available directly. Assuming that distinct folding states shall be enhanced, not only the order of force application (protocol #1 vs protocol #2) but also the choice of forces is crucial. Here, we demonstrate that selective modulation of the occupancies of a set of states, is directly possible by adjusting one force in the cycle, in this case *F*_high_ (Fig. 4h). *F*_high_ affects particularly the transition f_L_ → U, and, by extension, modulates the relative populations of states f_L_ and U. Lowering *F*_high_ from 45 to 33 to 30 to 27 pN increased the occupancy of f_L_ compared to U, thus the state f_L_ was relatively enhanced. Such an increased occupancy can be seen as a specific preparation of a distinct state in a multi-domain protein for further studies, e.g. distinct interaction studies with ligands or chaperones and the effects of those on relative occupancies. From a thermodynamics point of view, we would like to point out that these occupancies should not be interpreted as relative energetic stabilities, but require a full underlying kinetic model.

## VI. Conclusion

In this study we investigate the effects of periodic square functions of force on folding dynamics of (multidomain) proteins. We show that force spectroscopy models together with a master equation and *Monte Carlo* simulations are sufficient to reproduce complex experimental data, applied to a two states folder (*λ*_*YA*_), a modular multidomain protein (*L*_*x*8_) and a hybrid (λ_YA_-L) multidomain protein. The discrete periodic force protocols allow for quantitative extraction of stabilities and for a modulation of state occupancies. In contrast to earlier continuous, periodic force transitions, the discrete transitions allow the application of well-defined constant-force models [10] to extract stabilities. Regarding multidomain proteins, assuming independent domains, such periodic forces were able to enhance distinct intermediate state which was hard or impossible with traditional force-spectroscopy approaches [22]. We further show its applicability by dissecting states which would be normally entangled and modulating the occupancy of target states (selective modulation). Both periodicity and exact force levels provide multiple tunable parameters to prepare distinct folding states, which could be crucial to dissect large proteins and their folding pathways as well as to study misfolding pathways in presence or absence of interaction partners. While prior knowledge of underlying kinetic parameters is of advantage to adjust the periodic force cycle, it is not required and a *de novo* cycle optimization within an experiment is likely. We anticipate periodic forces enable expansion of understanding the folding intricacies of large complex proteins or folded RNA by accelerating slow transitions and slowing fast transitions, thus, by extension, shifting transition kinetics into the time window of observation.

## VII. Methods

### A. Cloning and purification

Subcloning and protein purification detailed protocols can be found in Supplementary Note 12. The plasmid encoding for (protein L)_x8_ was a kind gift from Julio Fernandez, as well as the empty magnetic tweezers vector haloTag-(spy0128)_x2_. We sub-cloned haloTag-*λ*_*YA*_-(spy0128)_x2_ and haloTag-*λ*_*YA*_-L-(spy0128)_x2_ using standard restriction-ligation and Gibson assembly protocols, respectively. Proteins were expressed in BL21 cells and purified using Ni-NTA affinity followed by size exclusion chromatography. Proteins were subsequently in vitro biotinylated.

### B. Magnetic Tweezers methods

A more detailed description of the instrument, surface preparation and force-calibration can be found in the methods and Supplementary Notes 1 and 2 of [10]. In brief: We used a custom-built setup following earlier designs [28] on top of a Nikon Eclipse Ti inverted microscope equipped with a 100x S-fluor NA1.3 oil immersion objective (Nikon), illuminated with a Cool-LED pE100 white lamp and controlled with a C-focus piezo position system (Madcity labs) for bead tracking using image-analysis based methods implemented in Visual Studio Pro 2017. To control applied force we used two lines of neodymium grade N52 magnets (D33, K&U Magnetics) in vertical configuration, with fast (0.7 m/s) and precise (150 nm resolution) z-control by means of a Linear voice-coil (LFA-2010, Equipment solutions). Feedback systems (PID) for smooth movement of the voice-coil are implemented though a DAQ card (NI USB-6341, National Instruments). Applied force was calibrated by means of the well-studied force-extension characteristics of protein L as described earlier [10,28]. Based on the WLC extension expectations and a mono-exponential decay of force with magnets distance to the surface we fit the parameters that determine applied force at different magnets position between 0.12 and 100 pN for M-270 superparamagnetic beads. Surface silanization was used to attach O4-ligands (targeting the HaloTag domain) to the surface and to immobilize the amino-functionalized reference beads for magnetic tweezers drift correction. Every measurement was performed at identical conditions with freshly titrated and filtered PBS-BSA 1%, 10 mM ascorbic acid, pH 7.4; and streptavidin coated M-270 magnetic beads to apply force in single molecule tethers [10]. Single molecule fingerprinting is done by means of an expected difference in contour length Δ*L*_*fu*_ (*λ*_6− 85_) = 26.5 *nm* with reversible kinetics at 6-10 pN, and successive Δ*L*_*fu*_ (*L*) = 18.6 *nm* unfolding steps for protein L; we consistently use a persistence length of 0.5 nm as measured before in PBS-BSA [22] and a room temperature of *T* = 297.15 *K* for contour length estimations.

### C. Data analysis

All methods where implemented in IgorPro 10 (Wavemetrics), except for Langevin dynamics simulations which were implemented also in MATLAB R2023b available in the GitHub repository https://github.com/Hartmann-lib/Energy_Potential_Diffusion_Simulation. A detailed description of our kinetics analysis pipeline is described in Supplementary note 4 of [10]. To analyze occupancies, we used gaussian fitting based estimations of the area under the curve of each sub-population within a steady-state histogram typically with at least hundred transitions. States segmentation and dwell times analysis was achieved by means of thresholding algorithms that facilitate a sensitive and intuitive way to measure dwell-times. Instead of a single threshold for a two-states protein, we used two thresholds acting depending on the current state of the molecule. For multidomain proteins we used the same logic but with a larger number of thresholds, which works well when the unfolding pathway is known (as was our case). For rates fitting, instead of fitting exponential models to the dwell times distribution, we fitted

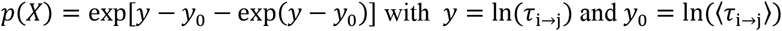

To the logarithm of dwell times distribution which allows to identify more easily artifacts from analysis or multiexponential kinetics. *Monte Carlo* simulations, numerical solutions of the master equation and Langevin dynamics methods are explained in detail in Supplementary Note 4, accompanied with pseudo-codes for implementation in Supplementary Note 11.

## Data availability

Data available upon request, with exemplary traces uploaded as supplementary raw data.

## Acknowledgements

We thank all members of the Schlierf lab for lively discussions during the development of this project. We thank the Julio Fernandez lab and Mordjane Boukhet for support with the initial set up of the instrument and appreciate insightful discussions with Felix Ritort. This project was partially supported by the Deutsche Forschungsgemeinschaft (DFG, German Research Foundation) under Germany’s Excellence Strategy – EXC-2068 – 390729961-Cluster of Excellence Physics of Life of TU Dresden and TU Dresden core funds (to M.S.). This research was funded by TU Dresden, DFG projects SCHL1896/6-1 and SCHL1896/9-1 (to. M.S.). This project is co-funded by the European Union and co-financed from tax revenues on the basis of the budget adopted by the Saxon State Parliament for the project aCOS (to A.H. and M.S.).

## Author contributions

C.A.Q.-C. and M.S. conceptualized the study, C.A.Q.-C., A.H. and M.S. devised and validated the methodology, C.A.Q.-C. performed the investigation and formal analysis, A.H. and M.S. supervised the project from investigation to writing, C.A.Q.-C. wrote the original draft and all other authors reviewed and edited it. C.A.Q.-C. and M.S. acquired funding for the project.

