## Supplemental Materials for "Complex folding kinetics of multi-domain proteins disentangled with periodic forces by magnetic tweezers"

**Content:**

**Supplementary Note 1.** Summary of terminology.

**Supplementary Note 2.** Force spectroscopy models.

**Supplementary Note 3.** Detailed balance from Kramers' theory.

**Supplementary Note 4.** Modelling kinetics under periodic forces.

**Supplementary Note 5.** Landscape features of $\lambda_{YA}$.

**Supplementary Note 6.** Generating observed vs expected stabilities plot.

**Supplementary Note 7.** Summary of the effects of periodic forces in two-states folding.

**Supplementary Note 8.** Measured kinetics of protein L.

**Supplementary Note 9.** Effects of a finite observation time in first passage times.

**Supplementary Note 10.** Kinetics of $\lambda_{YA} - L$ hybrid.

**Supplementary Note 11.** Pseudo-codes for simulations.

**Supplementary Note 12.** Sample preparation: Sub-cloning and protein purification.

**Supplementary Note 1. Summary of terminology.**

Here we list some of the terms and definitions we used along the main text. Please have in mind that some terms are found often in literature but lack a consensus, so they are not very precisely defined.

**Recurrent variables:**

- $\mathbf{i}, \mathbf{j}$ **:** referring to an arbitrary state i or j.
- $\mathbf{f}, \mathbf{u}, \mathbf{t}$ **:** folded, unfolded and transition states, respectively.
- $\boldsymbol{\Delta X}_{\mathbf{ij}}$ **:** Difference in end-to-end extension between states i and j.
- $\boldsymbol{\Delta G}_{\mathbf{ij}}$ **:** Difference in free energy between states i and j.
- $\mathbf{P_i}$ **:** occupancy of state i (as defined below).
- $\mathbf{k_{i \to j}}$ **:** transition rate of a molecule from state i to state j.
- $\boldsymbol{F}$ **:** Force.
- $\boldsymbol{F_{low}}, \boldsymbol{F_{mid}}, \boldsymbol{F_{high}}$ **:** In a periodic-force cycle, the lowest, middle and highest applied force respectively.
- $\mathbf{T_{low}}, \boldsymbol{T_{mid}}, \boldsymbol{T_{high}}$ **:** In a periodic-force cycle, the time spent in the lowest, middle and highest applied force, respectively.

**System in thermodynamic Equilibrium:** A system with no macroscopic flows (under steady state), time-reversible, and where we can apply the formalism of classical thermal physics (e.g. Boltzmann statistics for a closed system in contact with a heat reservoir). In force spectroscopy typically a constant force or constant extension measurement. We refer to **Out-of-equilibrium** when any of these conditions do not hold. In force spectroscopy, typically a force-jump, constant velocity or force-ramp where conditions (applied force) are changing.

**Quasi-equilibrium:** Simple terminology we used here to refer to out-of-equilibrium systems under steady state, where we cannot apply Boltzmann statistics. It is not particularly different from nonequilibrium steady states (NESSs) as discussed in literature [1], but it help us to keep language short and fluid. Off course, NESSs had been applied to a broader spectrum of phenomena.

**Steady state or stationary state observable:** Referring to an observable $Y_{obs}$ that do not change with time, $\partial_t Y_{obs} = 0$ . In this paper we often use occupancy, and label steady state occupancies with a "ss" superscript. We used the term as well for systems ("system at steady state") referring to their macroscopic observables.

**Elastic properties:** Often used in force spectroscopy, refers to the parameters that define how molecules respond to a shift in force. E.g. For a two state molecule with folded, unfolded and transition state in between, the difference in end-to-end extensions $\Delta X_{fu}(F, p, \Delta L_{fu}), \Delta X_{ft}(F, p, \Delta L_{ft})$, and $\Delta X_{tu}(F, p, \Delta L_{tu})$, given that they can be modelled as entropic springs.

**Mechanical midpoint or midpoint force:** The constant force at which it is equally likely to observe the molecule in the folded or unfolded state. $k_{f \to u}(F_m) = k_{u \to f}(F_m)$

**State Occupancy:** The probability of finding a molecule in a particular state, in this study typically within a time measurement, but often also defined within a number of iterations/molecules.

**Stability:** We use the term here strictly for $ln\left(\frac{P_i}{P_j}\right)$, in units of $k_B T$ for equilibrium conditions. We use stability instead of difference in free energy given that in out-of-equilibrium we cannot speak of stability in terms as a free energy difference.

**Supplementary Note 2. Force spectroscopy models.**

Polymer physics models

Whenever necessary, we estimate differences in end-to-end extensions by means of polymer physics model, namely the Marko-Siggia Worm-Like-Chain (WLC) interpolation formula [8,9]

$$F_{\mathrm{WLC}} = \frac{k_B T}{p}\left[\frac{1}{4}\left(1-\frac{x}{L}\right)^{-2} - \frac{1}{4} + \frac{x}{L}\right] \quad \text{(Eq. S2.1)}$$

With persistence length of 0.5 nm [10] . We estimate contour lengths using 0.365 nm / aa as measured earlier [11]. To extract extensions we shape Eq. S1 into a cubic polynomial together with Cardano's formula [12]. More details on how to account for spacer contributions and estimate energetics can be found in the supplementary section of [13].

Effects of force in stability

To extract the midpoint force ($F_m$) and the characteristic force response factor $\gamma$ of a two-states folding molecule we described occupancy vs force plots with a sigmoidal distribution, for instance for the folded state occupancy,

$$P_{\mathrm{f}}(F) = 1 - \frac{1}{1+e^{-\frac{F-F_{\mathrm{m}}}{\gamma}}} \quad \text{(Eq. S2.2)}$$

For a larger amount of states in equilibrium you can find a more involved description in the supplementary section of [14].

Effects of force in kinetics

For a the two-states system we used Bell-Evans model [15],

$$\mathrm{k}_{\mathrm{u}\to\mathrm{f}}(F) = k^0_{\mathrm{u}\to\mathrm{f}} \exp[-\beta(F\Delta X_{\mathrm{tu}})] \quad \text{(Eq. S2.3)}$$

$$\mathrm{k}_{\mathrm{f}\to\mathrm{u}}(F) = k^0_{\mathrm{f}\to\mathrm{u}} \exp[\beta(F\Delta X_{\mathrm{ft}})] \quad \text{(Eq. S2.4)}$$

And/or the MTS-model (Magnetic Tweezes model with Spacer correction as implemented in [13]),

$$k_{\mathrm{u}\to\mathrm{f}}(F) = k^0_{\mathrm{u}\to\mathrm{f}} \exp\left\{\beta\left[\int_{X^{\mathrm{p}}_{\mathrm{t}}}^{X^{\mathrm{p}}_{\mathrm{u}}} F^{\mathrm{p}}_{WLC}dx - \int_{X^{\mathrm{s}}_{t}}^{X^{\mathrm{s}}_{\mathrm{u}}} F^{s}_{FJC}dx - F\Delta X^{\mathrm{p+s}}_{\mathrm{tu}}(F)\right]\right\} \quad \text{(Eq. S2.5)}$$

$$k_{\mathrm{f}\to\mathrm{u}}(F) = k^0_{\mathrm{f}\to\mathrm{u}} \exp\left\{\beta\left[F\Delta X^{\mathrm{p+s}}_{\mathrm{ft}}(F) - \int_{X^{\mathrm{p}}_{\mathrm{f}}}^{X^{\mathrm{p}}_{\mathrm{t}}} F^{\mathrm{p}}_{\mathrm{WLC}}dx - \int_{X^{\mathrm{s}}_{t}}^{X^{\mathrm{s}}_{\mathrm{f}}} F^{\mathrm{s}}_{\mathrm{FJC}}dx\right]\right\} \quad \text{(Eq. S2.6)}$$

With superscripts p for protein contributions, and s for spacer contributions. Typically for simplicity assuming $\mathrm{X}^{s}_{\mathrm{t}} = 0$. For zero-force extrapolations large discrepancies arise for both models, but in our periodic force protocols, the range of extrapolations was typically very small (at most 2 pN away from the midpoint force), then they overlapped. Due to this for many applications even only using the Bell-Evans model would be sufficient. As a quality control at constant force we used detailed balance, as in equation (1) in the main text.

**Supplementary Note 3. Detailed balance from Kramers' theory.**

The equilibrium average free energy of the folded state is defined as:

$$\langle G_{\mathrm{f}} \rangle = -\frac{1}{\beta} \cdot ln(Z_{\mathrm{f}}) \qquad \text{(S3.1)}$$

with $\beta = 1/(k_B T)$ and $Z_F$ being the partition function. The partition function is defined as:

$$Z_{\mathrm{f}} = \int e^{-\beta \cdot G(X)} dx \qquad \text{(S3.2)}$$

Approximating a harmonic potential for the energy well of the folded state:

$$G(X) = \frac{\kappa_{\mathrm{f}}}{2}(X - X_{\mathrm{f}})^2 + G^{\mathrm{min}}(X_{\mathrm{f}}) \qquad \text{(S3.3)}$$

with curvature $\kappa_F$ and center position $x_F$, the partition function can be derived to:

$$Z_{\mathrm{f}} = e^{-\beta \cdot G^{\mathrm{min}}(X_{\mathrm{f}})} \int_{-\infty}^{\infty} e^{-\frac{\beta k_{\mathrm{f}}}{2}(X - X_{\mathrm{f}})^2} dx = \sqrt{\frac{2\pi}{\beta \kappa_{\mathrm{f}}}} \cdot e^{-\beta \cdot G^{\mathrm{min}}(X_{\mathrm{f}})} \qquad \text{(S3.4)}$$

Inserting this value in the first equation we obtain:

$$\langle G_{\mathrm{f}} \rangle = G^{\mathrm{min}}(X_{\mathrm{f}}) - \frac{1}{2\beta} \cdot ln\left(\frac{2\pi}{\beta \kappa_{\mathrm{f}}}\right) \qquad \text{(S3.5)}$$

for the curvature biased equilibrium free energy of the folded state. Applying this quantity and the similar equation for the unfolded state to the reaction equilibrium we derive:

$$\underbrace{K_{\mathrm{Eq}} = exp(-\beta \cdot \langle \Delta G_{\mathrm{fu}} \rangle)}_{\text{detailed balance}} =$$

$$\underbrace{\left(\frac{\kappa_{\mathrm{f}}}{\kappa_{\mathrm{u}}}\right)^2 exp\left(-\beta \cdot (G^{\mathrm{min}}(x_{\mathrm{u}}) - G^{\mathrm{min}}(x_{\mathrm{f}}))\right)}_{\text{Kramers}'\text{theory}} \qquad \text{(S3.6)}$$

This proves that the equilibrium constant of detailed balance as in the main text and Kramers' theory are the same differing only by the free energies $G_j$ and $G(x_j)$, which in the first case includes the curvature contribution and in the second case not.

**Supplementary Note 4. Modelling kinetics under periodic forces.**

Note: Please find IGOR Pro 10 pseudo-codes that accompany these methods in supplementary note 10.

As described in the main text, steady state occupancies at thermodynamic equilibrium follow detailed balance (Eq. (1)); kinetics (at equilibrium or out of equilibrium), follow Kramers' rate theory (Eq. (2)); and SMFS models describe the effects of force in kinetics (as summarized in supplementary note 2). For a given set of states $S_N = \{1,2,3,\dots,N\}$ the evolution of a system can be described by means of the master equation (Eq. 3). For instance, for a two states system ( $S_2 = \{1,2\}$ ) at constant force (wich means constant rates), the evolution of occupancies follows

$$\partial_t P_1(t) = -k_{1\to2}(F)\cdot P_1(t) + k_{2\to1}(F)\cdot P_2(t) \qquad \text{(S4.1)}$$

$$\partial_t P_2(t) = k_{1\to2}(F)\cdot P_1(t) - k_{2\to1}(F)\cdot P_2(t) \qquad \text{(S4.2)}$$

as described elsewhere [2,3]. These set of equations have an analytical solution of the form

$$P_1(t) = [P_1(t=0) - P_1^{ss}]\exp[-(k_{2\to1} + k_{1\to2})\,t] + P_1^{ss} \qquad \text{(S4.3a)}$$

$$P_2(t) = 1 - P_1(t) \qquad \text{(S4.3b)}$$

With $P_i(t=0)$ the initial conditions and $P_i^{ss}$ steady state occupancies. Well then, at periodic forces one would have to consider now two sets of rates, $k_{i\to j}(F^{high})$ and $k_{i\to j}(F^{low})$, which determine the evolution of the system at each given force. Border conditions must be considered to ensure continuity of $P_i(t)$, which is the reason why rates alone are not enough to estimate steady state occupancies. For certain force protocols (e.g. $F^{high} \gg F^{low} \sim F_m$ as detailed in supplementary note 5), the average occupancy over a periodic force cycle $\langle P_i(t)\rangle_T$ can be solved analytically, but very often (and specially for complex molecules) it is best to perform numerical solutions or simulations. To solve Eqs. 5 numerically we discretize time as follows,

$$P_1(t+\Delta t) = (1 - k_{1\to2}\Delta t)P_1(t) + (k_{2\to1}\Delta t)P_2(t) + O(\Delta t^2) \text{ (S4.4a)}$$

$$P_2(t+\Delta t) = (k_{1\to2}\Delta t)P_1(t) + (1 - k_{2\to1}\Delta t)P_2(t) + O(\Delta t^2) \text{ (S4.4b)}$$

With $k_{i\to j}\Delta t \ll 1$. In matrix notation the master equation can be written as $\vec{p}(t+\Delta t) = \mathbf{A}\vec{p}(t)$ with $\mathbf{A}$ the transition probability matrix. Discarding $O(\Delta t^2)$ terms, the elements of $\mathbf{A}$ become $A_{ij} = p_{ij} \approx k_{i\to j}dt,$ and $A_{ii} \approx 1 - \sum_{i\neq j}(k_{i\to j}\Delta t)$. Matrix notation becomes useful not only as it provides an algorithm to solve numerically the evolution of the master equation but it can be as well expanded to higher dimensional sets of states $S_N = \{1,2,3,\dots,N\}$. Moreover, at equilibrium it provides a straightforward way to estimate steady states, which can be easily achieved by finding the eigen vector that corresponds to eigen value 1. Effective rates can be easily estimated for square periodic forces,

$$\langle k_{i\to j}\rangle = \frac{k_{i\to j}(F_{low})T_{low} + k_{i\to j}(F_{high})T_{high}}{T_{low} + T_{high}}, \qquad \text{(S4.5)}$$

Which had been proposed as a way to estimate steady state occupancies [4]. But as mentioned earlier we think it is best to solve steady states numerically as exemplified in Fig. 3d. Effective rates do not work for every force protocol ($T_{low}, T_{high}, F_{low}, F_{high}$ ), an example being a very low $F_{low}$ can arbitrarily increase the folding rate.

*Monte carlo* simulations [5] help to simulate single molecule trajectories and hence are better suit to our type of data, and overall to describe phenomena that not only regard state occupancies but directly residence time distributions [6]. For this a simple algorithm starts with a transition probability following Poisson statistics, which again discarding terms of the order $O(\Delta t^2)$ ends up in the same transition probability matrix $\mathbf{A}$ as described earlier. Now instead of solving state occupancies, we observe the evolution of a random variable $s_t \in S_N$, generating a finite-state, discrete-time Markov chain where the probability to be in a state at time t, only depends on the previous state at time $t-1$,

$$\Pr\{s_t | s_{t-1}, s_{t-2}, \dots, s_0\} = \Pr\{s_t | s_{t-1}\} \qquad \text{(S4.6)}$$

Once a chain of states had been simulated, we converted our states to the space of extensions by using a three-dimensional matrix $B_{Fil}$ which at a given force (Force index F), stores the mean WLC extension (index $l = 0$) and standard deviation (index $l = 1$) for each state $i = s_t$. A simple workflow to estimate hidden Markov Chains for molecules with unknown kinetics using available open source programs [7] is suggested in supplementary note 13.

Finally, to further validate the relation between kinetics-predicted and observed stabilities we used overdamped Langevin dynamics simulations as implemented in [21]. We used the Euler-Mayurama integration scheme,

$$x(t+\Delta t) = x(t) - \gamma_x^{-1} \Delta t \partial_x G(x, F(t)) + \sqrt{2k_B T \gamma_x^{-1} \Delta t} R_X(t) \qquad \text{(S4.7)}$$

With $x(t)$ the simulated end-to-end distance of our molecules; $\gamma_x$ the friction coefficient; $R_x$ uncorrelated Gaussian random numbers with mean zero and unit variance; and $G(x, F(t))$ a time dependent energy landscape oscillating together with an applied force $F(t) = F^{high/low}$. After a Langevin dynamics simulation we treated the data in the same way as the experimental data following deriving kinetics, parameter models, etc.

### Supplementary Note 5. Landscape features of $\lambda_{YA}$

**Table S1.** Energy landscape of $\lambda_{YA}$.[a]

| Energy barriers[b] | | | | |
|---|---|---|---|---|
| **Model** | $k^0_{u\to f}$ [s$^{-1}$] | $k^0_{f\to u}$ [s$^{-1}$] | $\Delta G^0_{U-TS}$ [k$_B$T] | $\Delta G^0_{F-TS}$ [k$_B$T] |
| **Bell-Evans model** | (8.9±1.1)·10$^5$ | (4.1±2.2)·10$^{-4}$ | 5.6±0.5 | 27.1±0.1 |
| **MTS-model** | (4.2±2.5) ·10$^5$ | 0.8±0.4 | 13.2±0.5 | 26.4±0.6 |
| Elastic properties | | | | |
| **Model** | $\Delta X_{tu}(f_m)$ [nm] | $\Delta X_{ft}(f_m)$ [nm] | $\Delta L_{tu}$[nm] | $\Delta L_{ft}$[nm] |
| **Bell-Evans model** | 6.7±0.4 | 4.1±0.3 | - | - |
| **MTSS-model** | 7.6±0.9 | 5.5±0.6 | 17.3±1.9 | 12.5±1.3 |

[a] Error bars correspond to the SE of n=9 independent tethers each fitted separately with the Levenberg-Marquardt algorithm in igor pro.

[b] Barrier size estimations are done by assuming a Kramers' rate pre-factor of $1/k^0_{u\to f}$ of $1.5\ \mu s$ as measured for $k^0_{u\to f}\exp(-\beta\Delta G_{ut})$ for $\lambda_{6\text{-}85}$.

**Supplementary Note 6. Generating observed vs expected stabilities plot.**

Constant force data:

1. Measure $ln(P_u^{ss}/P_f^{ss})$ from raw extension histograms.
   a. Make a histogram of extensions of a trace with hundreds of transitions.
   b. Use multi-fitting to estimate $P_i^{ss}$ as in supplementary note 2.
2. Measure $ln(P_u^{ss}/P_f^{ss})$ as expected from kinetics.
   a. Measure dwell times histogram with hundreds of transitions as explained in supplementary note 2.
   b. Use the measured rates as an input for Montecarlo simulation. Master equation solvers also work, but care has to be taken on letting the system to equilibrate for a long enough time.
   c. Then find $ln(P_u^{ss}/P_f^{ss})$.

Periodic force data:

1. Measure $ln(P_u^{ss}/P_f^{ss})$ from raw extension histograms.
   a. Make a histogram of extensions of a trace with hundreds of transitions.
   b. Use multi-fitting to estimate $P_i^{ss}$ as in supplementary note 2.
2. Measure $ln(P_u^{ss}/P_f^{ss})$ as expected from kinetics.
   a. Using force spectroscopy models, estimate all four rates that determine $k_{i\to j}(F_{high})$ and $k_{i\to j}(F_{low})$, either by interpolation or by extrapolation.
   b. Use the measured rates as an input for *Monte carlo* simulations. Master equation solvers also work, but care has to be taken on letting the system to equilibrate for a long enough time.
   c. Then find $ln(P_u^{ss}/P_f^{ss})$.

**Supplementary Note 7. Summary of the effects of periodic forces in two-states folding.**

Let's define an asymmetry index as $\Lambda = \Delta X_{tu}/\Delta X_{ft}$, with a value of $\Lambda = 1$ for a perfectly symmetric potential of mean force (PMF). For instance, $\Lambda \sim 1.36$ for $\lambda_{YA}$ while $\Lambda \sim 25$ for protein L [19]. With this said here are a couple lessons we gained from studying the stabilities $\ln(P_u/P_f)$ of two-state folders, with the assumption of memoryless Markovian dynamics:

1. If the molecule PMF is symmetric, no stabilization effects of the folded or unfolded state are observed in force protocols surrounding the mechanical midpoint (i.e., $T_{high} = T_{low}$ and $F_{high/low} - F_m = 0$). Our experimental data corroborates this observation for a mildly asymmetric ($\Lambda = 1.63$) protein such as $\lambda_{YA}$, where only stabilizations of $\pm k_B T/2$ are observed.
2. If the protein is asymmetric, under protocols surrounding the mechanical mid-point (again i.e. $T_{high} = T_{low}$ and $F_{high/low} - F_m = 0$), you expect stabilization effects in a frequency and amplitude dependent manner, for a given landscape, typically at periods shorter than your molecular relaxation kinetics. You can induce $\frac{P_u}{P_f} = 1$ by setting $T_{high} > T_{low}$ or $F_{high} - F_m > F_{low} - F_m$, given that $\Lambda > 1$. As a remark, these protocols go beyond what is easily achievable with sinusoidal forces.
3. When taking into account force protocols with $T_{high} \neq T_{low}$ or $F_{high} - F_m \neq F_{low} - F_m$, the effects are various, and are better to be modelled for each particular case. Here a few examples which we think might be instrumental for technical developments and mechanobiology:
   a. $F_m - F_{low} \gg F_{high} - F_m \approx 0$. Which means going from a low force to the mechanical midpoint. Here the $T_{high}$ might act as a filtering parameter that cuts down slow transitions.
   b. $F_{high} - F_m \gg F_m - F_{low} \approx 0$. Which means going from the mechanical midpoint to a high force. A similar effect is expected but with $T_{low}$ as filter parameter. This protocol is particularly instrumental as a fitting protocol to get insights from kinetics from steady states analysis.

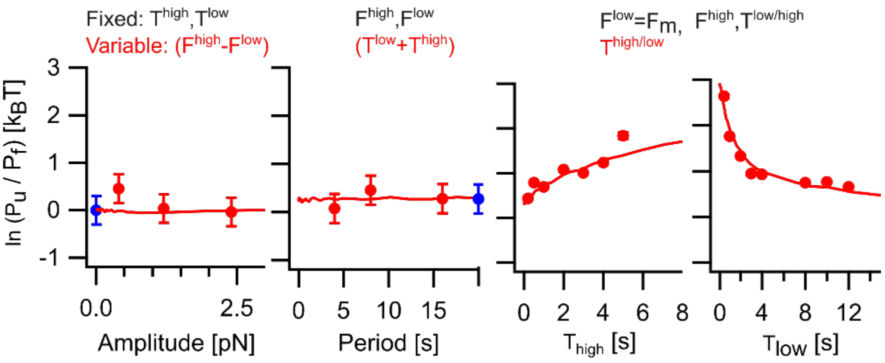


**Figure S1. Alternating forces produce stabilization effects according to a given force-protocol.** Here we show different variations of periodic force procols and their corresponding stabilization effects for $\lambda_{YA}$. Markers correspond to experimental data and solid lines to *Monte carlo* simulations. Blue marks correspond to constant force conditions.

4. Given that force is changing with time, it is crucial to understand that the kinetics of each molecule play a big role in these set of stabilization effects. Nevertheless, the exact role of kinetics in stability is in general not trivial, given that the condition $\langle P_u/Pf\rangle = \langle k_u\rangle/\langle k_f\rangle$ does not always hold. Overall, under sufficiently long periods of time, residence time distributions have multiexponential behaviors. Nevertheless, at periods close to molecular kinetics, deviations from multiexponential residence time distributions arise. We show here an exemplary residence time distribution with resonance peaks that deviates from multi-exponential dynamics.

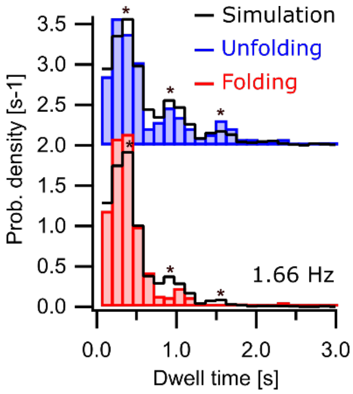


**Figure S2. Deviations from multiexponential kinetics for $\boldsymbol{\lambda_{YA}}$.** In this example we alternate forces at a frequency close to half the midpoint rates of $\lambda_{YA}$ (time matching condition), generating resonance peaks as described thoroughly earlier [6,20].

**Supplementary Note 8. Measured kinetics of protein L for *Monte Carlo* simulations.**

Qualitatively, literature values are enough to predict the state occupancies of each of the 8 states of $L_{x8}$ under periodic forces. But to gain quantitative agreement we measured kinetics for a given tether using the experimental data with the longest period, in our case of 32 seconds (16 s at $F_{\text{high}}$ and 16 seconds at $F_{\text{low}}$). Here then we extracted the dwell times using thresholding algorithms as described in main methods, measuring both, the dwell time of each state and the cumulative time that it takes from the moment we apply a high force (Figure S3 a). From the dwell time of each state, we fitted a function of the form

$$p(X) = \exp[y - y_0 - \exp(y - y_0)] \text{ with } y = \ln(\tau_{\text{i}\to\text{j}}) \text{ and } y_0 = \ln(\langle\tau_{\text{i}\to\text{j}}\rangle) \quad \text{(S13)}$$

Extracting then their dwell times $\langle\tau_{\text{i}\to\text{j}}\rangle$ (Figures S3b and c). Interestingly, scaling of $\langle\tau_{\text{i}\to\text{j}}\rangle$ does not scale up with degeneracy in a one-to-one fashion, which could be a consequence of linker effects from each of the unfolded subdomains. With this into account, we used for our *Monte Carlo* simulations the rate values

$$\begin{array}{ll} k_{0\to1} = 2.00\ s^{-1} & k_{4\to5} = 0.57\ s^{-1} \\ k_{1\to2} = 1.08\ s^{-1} & k_{5\to6} = 0.49\ s^{-1} \\ k_{2\to3} = 0.77\ s^{-1} & k_{6\to7} = 0.35\ s^{-1} \\ k_{3\to4} = 0.62\ s^{-1} & k_{7\to8} = 0.26\ s^{-1} \end{array}$$

together with a time step of 10 ms to estimate transition probabilities. We measured as well the mean extension and variance of each of the states under a given force from our experimental data and assigned them to our simulations in order to superimpose our simulated extension histograms with experimental ones. This can be done be means of a rank-3 tensor (three-dimensional matrix) with the first index to account for force, and the second and third indexes to save the means and variances of each state.

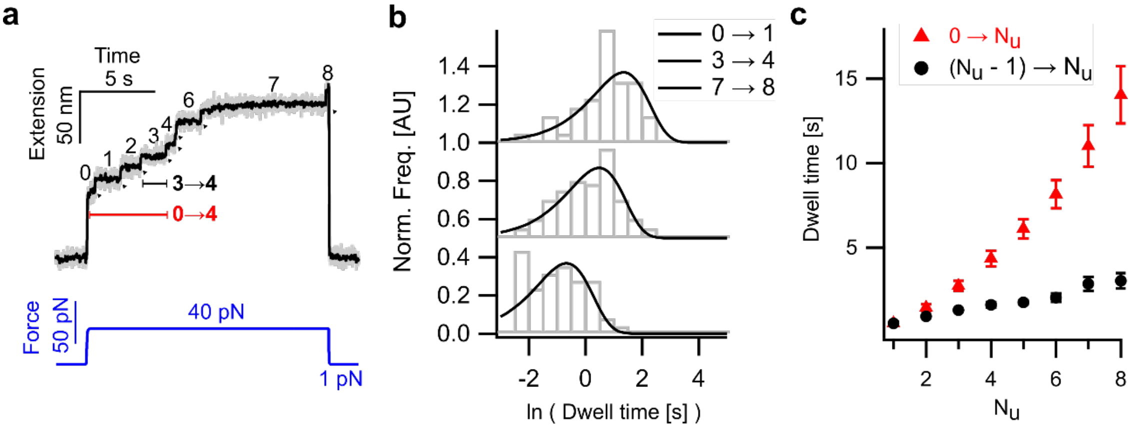


**Figure S3. Kinetics of Protein L unfolding. a)** Exemplary force jump with 9 states as labelled in the main text, Figure 3. We measured both the dwell time of each state (in black) and the cumulative dwell time to reach a given state from the moment force is applied (in red). **b)** exemplary residence time distributions for three of the 8 measured transitions and their respective fits. **c)** Fitted dwell times for all 8 possible transitions, both, per state transition and cumulative.

**Supplementary Note 9. Average first-passage time under a finite time window for $L_{x8}$.**

Residence times $\tau_i$ can be described by means of a Poisson probability distribution,

$$P(\tau_i) = \frac{1}{\langle\tau_i\rangle}\exp\left(-\frac{\tau_i}{\langle\tau_i\rangle}\right), \text{(S9.1)}$$

with mean and variance given by $\langle\tau_i\rangle$. In our $L_{x8}$ unfolding experiment, each individual subdomain is connected in series, hence it is safe to assume a constant applied force to every sub-domain individually displaying then single-exponential unfolding kinetics. Due to this, we can describe the first passage time distributions ($P(t_{i\to j})$) as well with Poisson statistics,

$$P(t_{i\to j}) = \frac{1}{\langle t_{i\to j}\rangle}\exp\left(-\frac{t_{i\to j}}{\langle t_{i\to j}\rangle}\right), \text{(S9.2)}$$

Which would not be in general the case for proteins with entangled subdomains with multi-exponential kinetics derived from more intricate unfolding pathways. $\langle t_{i\to j}\rangle$ is connected to the cumulative dwell time described in Fig 4b and in Supplementary note X. Now then, we can find the average $\langle t_{i\to j}\rangle$ over a finite time window $T_{obs}$, as

$$\langle t_{i\to j}\rangle\big|_{T_{obs}} = \int_0^{T_{obs}} tP(\tau_{i\to j})\, dt = \frac{\langle t_{i\to j}\rangle|_\infty}{1}\left\{1-\left[\frac{T_{obs}}{\langle t_{i\to j}\rangle|_\infty}+1\right]\exp\left(-\frac{T_{obs}}{\langle t_{i\to j}\rangle|_\infty}\right)\right\}, \text{(S9.3)}$$

which converges to $\langle t_{i\to j}\rangle\big|_\infty$ when $T_{obs}$ is sufficiently high. From here on, it is easy to plot different versions of this equation according to what is more meaningful to your data. For instance, we plotted $\langle t_{i\to j}\rangle\big|_{T_{obs}}/\langle t_{i\to j}\rangle\big|_\infty$ as a function of $T_{obs}/\langle t_{i\to j}\rangle\big|_\infty$ in the main text which is a way to normalize a $\langle t_{i\to j}\rangle\big|_{T_{obs}}$ vs $T_{obs}$ plot with respect to the steady state kinetics. A second meaningful value is $\langle t_{i\to j}\rangle\big|_{T_{obs}}/T_{obs}$ , which helps us investigate the probability of finding a $i \to j$ transition per unit time. Overall, for periodic forces $T_{obs}$ is constant, nevertheless in biology you will find often scenarios where $T_{obs}$ (or $T_{perturbation}$) is taken out from a probability distribution describing a different kind of cyclic process.

**Supplementary Note 10. Kinetics of $\lambda_{YA} - L$ hybrid.**

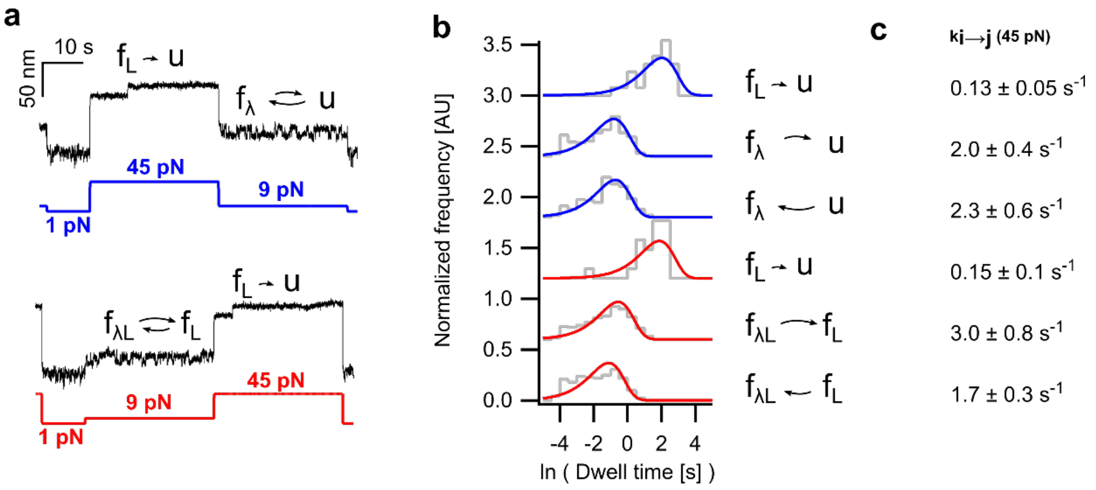


**Figure S4. Kinetics of the individual states within the hybrid $\lambda_{YA} - L$. a)** exemplary cycle (as in main figure 4) with three forces and five extension states with the observed kinetics highlighted above. **b)** residence time distribution for each observed type of transition fitted with its respective single-exponential. **c)** extracted rates for each of the transitions as schematized in a and b (±95% CI).

**Supplementary Note 11. Pseudo-codes for simulations.**

```
// this code simulates a single monte carlo step iterating up. This
facilitates uploading the rates in each step which helps then to define force
vs time.
Function SimulateDTMC(ku,kf,allpaths,step)
    Variable ku,kf,step
    wave allpaths

    Variable T = 1              // number of steps
    Variable n = 2               // Number of states
    Variable Nsim = 1           // Number of trajectories to simulate
    Variable i, j, z=step, step_time=(1/100) // time for each step in 10 ms.

    // Define transition matrix P
    Variable P12=ku*step_time, P21=kf*step_time


    Make/O/N=(n,n) Pr

    // 2-states
    Pr[][0] = {(1-P12), P12}
    Pr[][1] = {P21, (1-P21)}

    // Alternatively for 8-states (with respective variables initialization):
    Pr[][0] = {(1-P12), P12, 0, 0, 0, 0, 0, 0, 0}
    Pr[][1] = {P21, (1-P23-P21), P23, 0, 0, 0, 0, 0, 0}
    Pr[][2] = {0, P32, (1-P34-P32), P34, 0, 0, 0, 0, 0}
                    (…)
    Pr[][8] = {0, 0, 0, 0, 0, 0, 0, P98, (1-P98)}

    MatrixTranspose Pr

    // Simulate Nsim paths
    Variable current, r, cumP, nextState

    for (i = 0; i < Nsim; i += 1)
            current = allPaths[i][z - 1] // current state
            r = abs(enoise(1))  // uniform random number in [0,1)
            cumP = 0
            for (j = 0; j < n; j += 1)
                cumP += Pr[current][j]
                if (r < cumP)
                    nextState = j
                    break
                endif
            endfor
            allPaths[i][z] = nextState
    endfor

End
```

**Algorithm S11.1.** Pseudo-code for *Monte Carlo* simulations implemented in Igor Pro 10.

```
Function states2extensions()
    \\Load Markov chain MC. Markov chain[0][x] contains states vs time,
Markov chain[0][x] contains force-index vs time.

    wave MC

   \\Load B_fil, a three dimensional matrix with f an index to account for
force, i to account for state, l=0 mean extension and l=1 SD.

    wave B_fil
    Variable timepoints=numpnts(MC)/2

    Make /O /N=(timepoints) MC_extensions
    Make /O /N=(timepoints) MC_time
    Make /O /N=(timepoints) Variances_L

    MC_extensions=
B_fil[MC[0][x]][0][MC[1][x]]+gnoise(B_fil[MC[1][x]][1][MC[1][x]])

End
```

**Algorithm S11.2.** Pseudo-code to convert states into extensions implemented in Igor Pro 10.

```
Function SolveME(T,ku,kf, p0, probMatrix)
    Variable T, ku, kf      // Number of time steps T
    wave p0, probMatrix // po: initial conditions. probMatrix, outcomes.
    Variable n = 9              // Number of states
    Variable z, i, j
    Variable time_step=1/100 // 10 miliseconds.
    Variable P12=k12*time_step, P21=k21*time_step, (…)

    Make/O/N=(n,n) Pr

    Pr[][0] = {(1-P12), P12, 0, 0, 0, 0, 0, 0, 0}
    Pr[][1] = {P21, (1-P23-P21), P23, 0, 0, 0, 0, 0, 0}
    Pr[][2] = {0, P32, (1-P34-P32), P34, 0, 0, 0, 0, 0}
                        (…)
    Pr[][8] = {0, 0, 0, 0, 0, 0, 0, P98, (1-P98)}
    MatrixTranspose Pr
    // Numerically evolve: p_{t+1} = p_t * P
    Variable sum
    for (z = z0; z <= (T); z += 1)
        for (j = 0; j < n; j += 1)
            sum = 0
            for (i = 0; i < n; i += 1)
                sum += probMatrix[i][z - 1] * Pr[i][j]
            endfor
            probMatrix[j][z] = sum
        endfor
    endfor

End
```

**Algorithm S3.** Pseudo-code to solve numerically the master equation.

**Supplementary Note 12. Protocols for cloning and protein purification.**

Sub-cloning:

The (Protein L)$_{x8}$ construct for magnetic tweezers (Figure S1a) and the empty vector HaloTag-Spy0128$_{x2}$ where kindly provided by the members of the former Julio Fernandez Lab. Subcloning of the gene encoding for HaloTag-$\lambda_{6\text{-}85}$-Spy0128x2 (Figure 1a) is explained in detail in the supplementary section of [13], by means of restriction-ligation. Here we include an exemplary protocol as we used it for *haloTag-$\lambda_{YA}$-L-(spy0128)$_{x2}$* by means of Gibson Assembly, following standard protocols as provided by NEB.

Exemplary sub-cloning protocol: *haloTag-$\lambda_{YA}$-L-(spy0128)$_{x2}$*

Overall: Linearize a *halotag- $\lambda_{YA}$ -(spy0128)$_{x2}$* vector with BamHI, PCR to get an L insert with ~15nt overlapping sequence for Gibson Assembly, perform Gibson Assembly reaction, pick successful cloning with colony PCR and finally verify by sequencing.

Note: When designing primers for Gibson assembly with a vector linearized by restriction enzymes, particular care must be taken as one misplaced nucleotide can shift the ORF. SnapGene was used for design and cloning *in silico* before starting a cloning project.

For the Vector:

1. Incubate ON in LB with ampicilin *E. coli* XL10 cells containing *halotag- $\lambda_{YA}$ -(spy0128)$_{x2}$* cloned into pFN18a vector, as described in [13]. A Bam-HI restriction site remains in between $\lambda_{YA}$ and *(spy0128)$_{x2}$*.
2. Miniprep following provider instructions (QuickPure, Macherey-Nagel).
3. Digestion
   3.1. Mix 2 ug of plasmid, 5 µl Cutsmart buffer, 3 µl BamHI-HF [20 U] (New England Biolabs), and milliQ to reach a volume of 50 µl.
   3.2. Incubate 30 min at 37 °C.
   3.3. Clean up (Zymo Research).
4. Check DNA size with Electrophoresis (1.2% agarose gel).

For the Insert:

Design of primers following NEB instructions and verify cloning *in silico* using SnapGene.

5. PCR of insert with additional ~15 nt overlapping sequence for Gibson assembly.
   5.1. Fwd: 5' TTGCACGTGAAATTGGATCCATGGAAGAAGTAACTATTAAAGCTAATTT 3'
   Rvs: 5' TGAACTGTTGTAGCGGATCCGCTCCTAAATTTTATATTTAGAGTATATCCCT 3'
   5.2. PCR mix: 10 µl 5x Herculase Rxn Buffer, 5 µl dNTP [10 mM], 1.25 µl Fwd [10 µM], 1.25 µl Rvs [10 µM], 30 ng plasmid with $L_{x3}$, 1 µl Herculase II Fusion (Agilent) and nuclease free water up to 50 µl.
   5.3. Standard PCR protocol following provider recommendations for Herculase polymerase.
   5.4. Clean up and check insert size with Electrophoresis (1.2% agarose gel).

Gibson assembly:

6. Gibson assembly reaction and transformation.
   6.1. Mix 10 µl Gibson assembly master mix (2X), 100 ng vector, insert to a molar ratio 3:1 (insert:vector), and miliQ water up to 20 µl.
   6.2. Incubate 15 min at 50 °C and chill on ice.
7. Transform the ligation product into XL10 cells by heat shock transformation 30 seconds at 42 °C. Plate in LB with ampicillin.
8. Seek for *L* DNA containing colonies by colony PCR.
9. Validate positive results by miniprep followed by Sanger sequencing (Microsynth) (Figure S5).

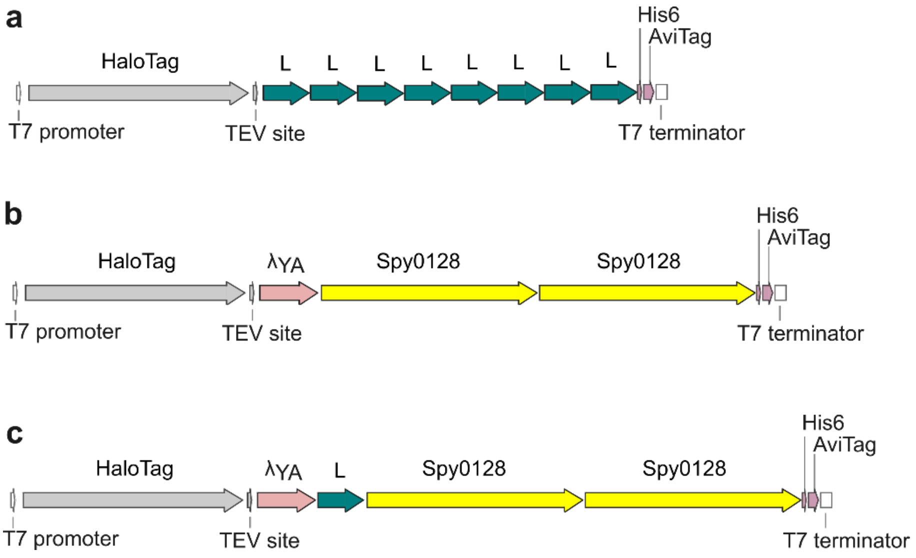


**Figure S5. Gene maps of our constructs. a)** HatoTag-(Protein L)$_{x8}$ construct with repetitive sequences of the B1 subdomain of protein L, **b)** recombinant HaloTag-$\lambda_{YA}$-Spy0128$_{x2}$ encoding gene and **c)** HatoTag-$\lambda_{YA}$-L-Spy0128$_{x2}$. All constructs are cloned in pFN18a. Sequences were all verified by Sanger sequencing (Microsynth).

<u>Protein purification (find more details in the supplementary section of [13])</u>

1. Overexpression, in TB or LB with ampicillin with 1mM IPTG for induction. Lysis buffer:10 mM HEPES, 300 mM NaCl, 1 mM DTT, 10% glycerol, pH 7.2 plus 1 protease inhibitor tablet (Roche) and 5 µl Benzonase endonuclease (>90% purity) (Merck-Millipore).

2. Standard Histrap purification, column HisTrap 5ml FF or HP (GE Healthcare). Binding buffer: 10 mM HEPES, 150 mM NaCl, 10% glycerol, 0.5 mM EDTA and 20mM Imidazole, filtered and degassed. Linear gradient elution with up to 0.5 M imidazole.

3. Size Exclusion Chromatography to increase purity. Column: Superdex 200 increase (10/300 GL) column (GE Healthcare). Size exclusion buffer: 10 mM HEPES, 150 mM NaCl, 10% glycerol and 0.5 mM EDTA, filtered and degassed.

4. Select for purity in each step with SDS-page (10% acrylamide gel).

5. Biotinylation (BirA500-RT kit, Avidity) followed by free biotin removal with Vivaspin6 (MWCO:30 kDa, Sartorius) protein concentrators.

6. Flash freeze and store. Storing buffer: 10 mM HEPES, 150 mM NaCl, and 10% glycerol; filtered and degassed.

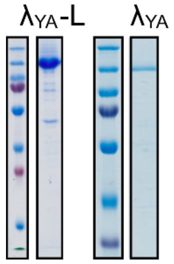


**Figure S6. Protein purification quality control.** SDS-page of pure HatoTag-$\lambda_{YA}$-L-Spy0128$_{x2}$ and HaloTag-$\lambda_{YA}$-(spy0128)x2 (Each next to their corresponding ladder). Page ruler plus pre-stained ladder (Thermo Scientific).
